# Convergent targeting of conserved regulatory networks during thermal evolution across *Saccharomyces*

**DOI:** 10.64898/2026.03.23.708575

**Authors:** Jennifer Molinet, Carolin Gierer, Pablo Villarreal, Rike Stelkens

## Abstract

Whether evolution follows predictable genetic paths across species remains a central question in evolutionary biology, particularly as rising temperatures reshape species distributions worldwide. Despite its importance, the genetic basis of thermal adaptation remains poorly understood across divergent species. Here, we use the yeast genus *Saccharomyces* as a comparative model to investigate how species with contrasting thermal niches adapt to rising temperatures. We combined experimental evolution under progressively increasing temperatures for up to ∼600 generations with whole-genome sequencing of 256 evolved genotypes, followed by transcriptomic, functional, and physiological analyses across eight species. Despite large differences in ancestral thermal tolerance and evolutionary outcomes, selection repeatedly targeted the same conserved regulatory pathways across species. Independent lineages accumulated *de novo* mutations in central growth and stress response networks, particularly in TORC1, PKA, and MAPK signaling pathways, revealing striking molecular convergence across species occupying distinct thermal environments. However, these shared genetic targets produced divergent transcriptional and physiological responses depending on species background, indicating that thermal adaptation primarily proceeds through rewiring of conserved regulatory hubs rather than changes in temperature-specific enzymes. Cold-tolerant species frequently lost mitochondrial DNA during thermal evolution, yet loss alone was insufficient to reproduce the adaptive thermal phenotypes of evolved populations. Together, our results show that adaptation to increasing temperature is driven by predictable changes in conserved regulatory networks, while species-specific constraints shape divergent phenotypic outcomes. These findings reveal both the predictability and contingency of evolutionary responses to rising temperature across species.

## INTRODUCTION

Thermal adaptation is a fundamental process that shapes the ecology, distribution, and evolution of organisms^1,2^. With global climate change driving more frequent and intense thermal extremes, understanding how organisms adapt at the molecular level is crucial for predicting their evolutionary trajectories and resilience^3,4^.

Species of the genus *Saccharomyces* provide a powerful model for studying thermal adaptation, given their broad diversity in thermal tolerance and ecological niches. The genus spans from cold-adapted species such as *S. eubayanus*, *S. arboricola*, and *S. kudriavzevii* to warm-adapted species such as *S. cerevisiae* and *S. paradoxus*^5–8^. This natural thermal diversity, combined with experimental tractability in the laboratory, enables controlled experimental evolution studies that allow direct observation of evolutionary changes over hundreds of generations^9–11^. Thermal performance curves (TPCs), which describe the relationship between temperature and organismal performance, provide a quantitative framework for linking evolutionary change to fitness-related traits and a common currency for comparing adaptive responses across species^12–15^. Previous work has shown that different *Saccharomyces* species evolve distinct TPCs when exposed to constant versus rising temperatures^5^. Here, we describe the molecular basis of these changes.

We previously evolved populations of eight *Saccharomyces* species under constant (25 °C) and progressively increasing temperature regimes, ranging from 25 to 40 °C, for up to 600 generations, to assess their evolutionary potential in adapting to future warming^5^. We found that TPCs varied significantly between species, revealing two main trajectories: i) Warm-tolerant species showed an increase in both optimum growth temperature and thermal tolerance, consistent with a “hotter is wider” evolutionary trajectory; ii) Cold-tolerant species on the other hand evolved larger thermal breadth and higher thermal limits, but suffered from reduced maximum performance overall, consistent with a generalist or “a jack of all temperatures is a master of none” trajectory^5^.

Despite these well-characterized phenotypic trajectories, the molecular mechanisms underlying thermal adaptation across species remain poorly understood. Thermal adaptation may involve changes in regulatory pathways, protein stability, membrane composition, and organellar function^7,16–21^. Although growing evidence indicates that temperature is a dominant selective force under ongoing climate warming^22^, it remains unclear whether adaptation to rising temperatures proceeds through predictable molecular solutions (convergent evolution) or is instead constrained by species-specific physiological and evolutionary histories (divergent evolution)^23,24^. While phenotypic adaptation is often repeatable^24^, growing evidence from experimental evolution and comparative genomic studies suggests that the underlying genetic basis can be highly contingent, with convergence frequently emerging at the level of molecular pathways rather than individual genes^23,25–27^. Consequently, we still lack an integrative understanding of how conserved cellular signaling pathways, organelle function, and organism performance interact to shape evolutionary trajectories under thermal stress. Especially mitochondrial function represents a putative yet understudied mechanism of thermal adaptation, given its central role in energy production, metabolic regulation, and stress responses, as well as its known sensitivity to temperature^28,29^.

Here, we use the budding yeast genus *Saccharomyces* as a comparative evolutionary model to address these gaps in our understanding of the molecular and regulatory basis of thermal adaptation. By combining whole-genome sequencing across eight species, functional analyses of conserved growth-stress signaling pathways, mitochondrial manipulations, and thermal performance assays, we ask whether climate warming drives convergent molecular evolution and how it manifests at the phenotypic and physiological levels. Our integrative approach reveals that while increasing temperature repeatedly targets conserved signaling hubs, adaptive responses are ultimately shaped by species-specific regulatory and energetic constraints, with important implications for predicting evolutionary responses to climate change.

## RESULTS

### Genetic changes associated with thermal evolution

To explore the genetic basis of TPCs evolution under increasing temperature conditions in eight *Saccharomyces* species, we analyzed *de novo* single-nucleotide polymorphisms (SNPs) across 256 evolved genotypes and their ancestral strains (**Table S1** and **S2**). The 256 evolved genotypes correspond to individual clones isolated at the end of experimental evolution, from populations across eight *Saccharomyces* species, including between one to three strains per species. Evolution was performed in four independent replicate populations per strain and temperature regime (constant and increasing), and two individual clones (genotypes) were sequenced per population. The genomes of each evolved genotype were compared to their corresponding ancestral strain, allowing for the identification of *de novo* mutations that arose during experimental evolution. Across all genomes, a total of 378 SNPs were identified, with variation in the number and genomic location across species, strains, genotypes, and temperature conditions (**Figure 1** and **S1**, **Table S2C**).

**Figure 1.**
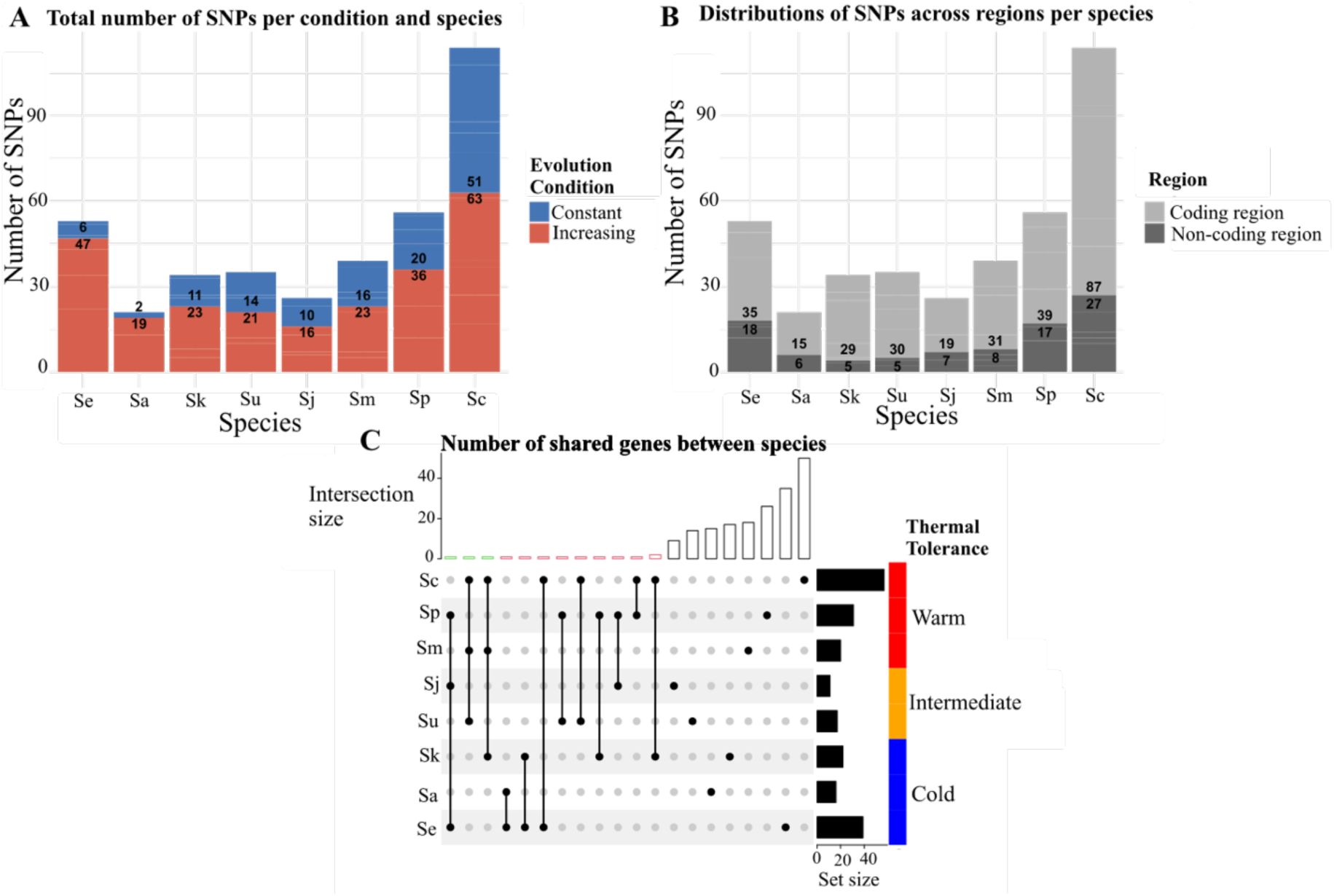
Genetic changes occurring in experimental evolution in constant vs. increasing temperatures in *Saccharomyces*. **(A)** Total number of *de novo* SNPs per species and experimental evolution condition (constant vs. increasing temperature). **(B)** Distribution of SNPs across coding and non-coding regions per species. **(C)** Parallel evolution across species revealed by independent mutations in shared genes, displayed as an UpSet plot. Species are grouped by thermal tolerance (blue = cold-tolerant, orange = intermediate, red = warm-tolerant). Set size bars show the number of mutations per species. Intersection size bars show the number of mutations shared by two or more species. The intersection matrix shows which species (lines) share mutations.

Genotypes evolved under increasing temperatures accumulated nearly twice as many *de novo* SNPs as those evolved under constant temperature (248 *vs*. 130 SNPs; **Figure 1A** and **S1A**), suggesting stronger selection pressures under increasing temperature regimes. SNPs were detected in both coding and non-coding regions, but mutations in coding sequences were on average three times more likely under both evolutionary conditions (**Figure 1B** and **S1B**). A subset of these mutations was synonymous or located in regulatory regions, suggesting possible functional consequences (**Figure 1B** and **S1B**).

To test whether mutational accumulation differed among species and temperature regimes, we fitted a generalized linear mixed-effects model using the number of *de novo* SNPs per evolved clone as the response variable, with species, evolutionary condition (constant vs. increasing temperature), and their interaction as fixed effects, and strain as a random effect. We detected significant effects of species (Wald test, p-value = 0.027) and evolutionary condition (Wald test, p-value < 0.001), as well as a strong species x condition interaction (Wald test, p-value < 0.001, **Table S2F**), indicating that the increase in the number of SNPs under thermal stress varies across species. These results suggest that evolution in increasing temperatures leads to faster accumulation of *de novo* SNPs than in constant temperature, but that the magnitude of genetic change is strongly modulated by the species’ genetic background.

To determine whether these mutations affect biological processes or signaling pathways, we performed GO term and KEGG pathway enrichment analyses. For genotypes evolved in increasing temperature, we found enrichment for genes related to cellular processes, positive regulation of cell growth, regulation of invasive growth in response to glucose limitation, and regulation of filamentous growth (**Table S2G**). Genotypes evolved in increasing temperature also showed an enriched KEGG pathway related to longevity regulation, involving the TORC1, PKA, and MAPK signaling pathways (**Table S2G**). Meanwhile, for genotypes evolved at constant temperature, we found enrichment for genes involved in the regulation of biological processes and primary metabolic processes, such as adenylate cyclase-modulating G protein-coupled receptor signaling pathways, and Ras protein signal transduction (**Table S2G**).

Notably, 13 genes repeatedly accumulated independent mutations across multiple strains and species under increasing temperature conditions, including species with different natural temperature tolerances (e.g., the warm-tolerant species *S. cerevisiae* and the cold-tolerant species *S. kudriavzevii*). These included *BMH1*, *SNF4*, *CYR1*, and *PRR2*, which are involved in cellular signaling pathways (e.g., TORC1 and MAPK). This indicates repeated targeting of the same functional components across genetically and ecologically vastly different species, suggesting convergent molecular evolution under thermal selection (**Figure 1C**). In comparison, genotypes evolved under constant temperature showed only four genes with independent mutations (shared by three species: *S. cerevisiae*, *S. uvarum*, and *S. kudriavzevii*; **Figure S1C**).

### Differential activation of signaling pathways

To evaluate the functional implications of the pathways detected by SNP enrichment analysis (TORC1, PKA, MAPK), we measured the expression of reported canonical target genes (TORC1: *RPS6A*, *CRF1*; PKA: *SSA4*, *TPK1*; MAPK: *SLT2*, *RLM1*)^21,30–34^ by qPCR in *S. cerevisiae* (the most warm-tolerant species) and *S. eubayanus* (the most cold-tolerant species)^5^, upon exposure to benign and high temperatures (**Figure 2 and S2**, **Table S3A**).

**Figure 2.**
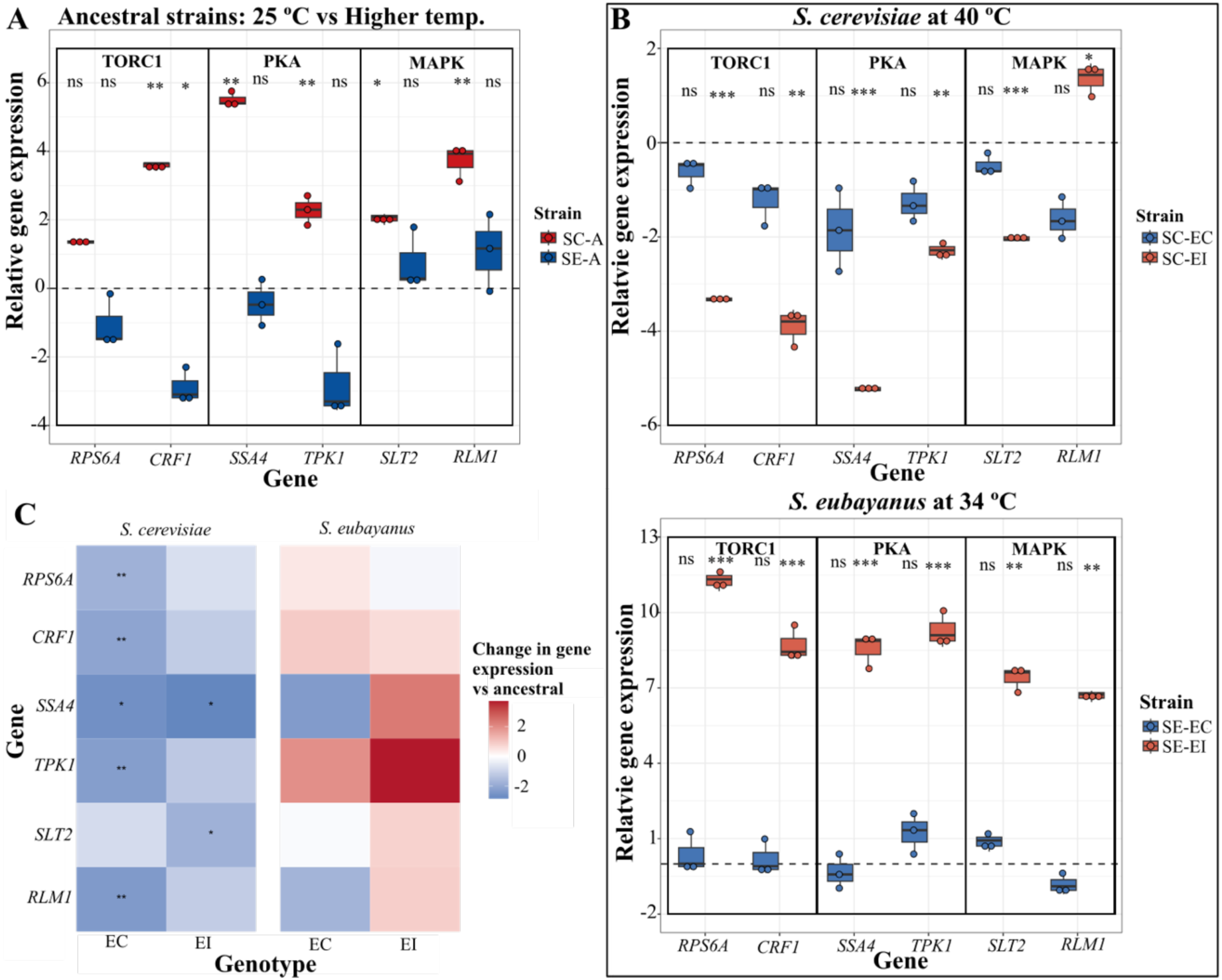
Functional validation of TORC1/PKA/MAPK outputs across species and temperature conditions. **(A)** Log_2_ normalized gene expression for ancestral *S. cerevisiae* (SC-A, red) and *S. eubayanus* (SE-A, blue) shown as the relative difference when exposed to 25 °C vs. the species-specific high temperature (40 °C for SC; 34 °C for SE). **(B)** Log_2_ normalized expression of evolved genotypes relative to their respective ancestral strains at high temperature (SC 40 °C, top; SE 34 °C, bottom). EC: genotype evolved at a constant 25 °C; EI: genotype evolved in increasing temperature. In (A) and (B), the dashed line marks no change. Plotted values are the average of three biological replicates. Asterisks denote significant differences in gene expression within species in (A) and evolved-ancestral differences in (B) (Welch’s t-test, * p < 0.05, ** p < 0.01, *** p < 0.001, ns = non-significant). **(C)** Heatmap of temperature effects on gene expression for evolved genotypes relative to their ancestors, computed as (expression at high temperature minus expression at 25 °C in EC or EI) minus (expression at high temperature minus expression in the ancestor). Blue indicates a smaller response to temperature (dampened plasticity); red indicates a greater response (enhanced plasticity). Asterisks mark significant temperature x evolved genotype interaction terms from OLS models with HC3 robust standard errors. Genes are grouped by pathway (TORC1: *RPS6A*, *CRF1*; PKA: *SSA4*, *TPK1*; MAPK: *SLT2*, *RLM1*).

First, we examined gene expression in the ancestral strains (before experimental evolution). At 25 °C, *S. cerevisiae* and *S. eubayanus* ancestors differed in several pathway readouts (**Figure S2A**). *S. cerevisiae* showed higher expression than *S. eubayanus* in *RPS6A*, *SLT2*, *SSA4*, and *RLM1*. *S. eubayanus* exceeded *S. cerevisiae* in *CRF1* and *TPK1* (Welch test, p-value < 0.05, **Table S3B**). This indicates species-specific baselines for TORC1/PKA/MAPK output under benign temperature conditions.

When ancestral strains were exposed to their species-specific high temperature (maximum temperature reached in experimental evolution: 40 °C for *S. cerevisiae*, 34 °C for *S. eubayanus*), *S. cerevisiae* showed robust induction of expression in *CRF1*, *RLM1*, *SLT2*, *SSA4*, and *TPK1* (log₂ ≈ +3.6, +3.7, +2.0, +5.5, +2.3; Welch tests, p < 0.05), but no significant change in *RPS6A*. In contrast, *S. eubayanus* showed significant repression of *CRF1* (log₂ ≈ −2.9; Welch tests, p < 0.05) but no statistically significant expression differences in the other genes (**Figure 2A**; **Table S3C**). A species x temperature interaction test (Ordinary Least Squares (OLS) model with HC3 robust standard errors) confirmed that the magnitude of the heat response (high temperature vs. 25 °C) differed between species for multiple genes (*CRF1*, *SSA4*, *TPK1*, *RPS6A*, p < 0.05, **Table S3C**). These patterns indicate a stronger transcriptional heat response in the warm-tolerant species (*S. cerevisiae*) and a limited or suppressed response in the cold-tolerant species (*S. eubayanus*), consistent with their contrasting thermal ecologies.

We next asked whether evolved genotypes (isolated from the endpoint of experimental evolution) diverged transcriptionally from their ancestors when assayed at high temperature (**Figure 2B**, **Table S3D**). For this, we compared genotypes evolved at constant (EC) and increasing temperature (EI) to their respective ancestors. In *S. cerevisiae*, EI genotypes showed consistent down-regulation of gene expression relative to their ancestor across all genes at 40 °C (*SSA4:* log₂ = −5.22; *RPS6A:* −3.32; *CRF1:* −3.89; *SLT2:* −2.03; *TPK1:* −2.30; Welch tests, p < 0.05) except for *RLM1,* where gene expression increased (log₂ = +1.37, p = 0.026, **Figure 2B**, **Table S3D**). In contrast, expression in *S. eubayanus* EI genotypes was strongly up-regulated relative to the ancestor for all genes at 34 °C, with log₂ normalized expression from +6.69 to +11.26 across *RLM1, SLT2, TPK1, SSA4, CRF1,* and *RPS6A* (all p ≤ 0.007, **Figure 2C**, **Table S3D**). Notably, we did not observe any differences between the constant-evolved genotypes (EC) and their ancestors when assayed at high temperature (**Table S3D,** Welch test, p > 0.05). This demonstrates that the major functional shift at high temperature is specific to EI genotypes. Together with the enrichment of mutations in TORC1/PKA/MAPK (**Figure 1**), these results suggest that evolution under increasing temperature elicits a stronger and directionally opposite remodeling of central signaling pathway activity in warm-vs. cold-tolerant yeast species, with repression in *S. cerevisiae* and hyperactivation in *S. eubayanus*.

We also measured the transcriptional response of the evolved genotypes (EI and EC) at 25 °C (**Figure S2B**, **Table S3E**). *S. cerevisiae* EI genotypes did not show any expression differences to their ancestor, whereas *S. eubayanus* EI genotypes showed widespread constitutive up-regulation (Welch test, p < 0.05). Thus, in *S. eubayanus*, the evolved shift was observable already at 25 °C, whereas in *S. cerevisiae* this manifested only at higher temperatures.

For each type of genotype (ancestral, EC, EI), we calculated the temperature effect as the change in expression between the high temperature and 25 °C (**Figure S2C**). We then tested whether evolution altered thermal regulatory plasticity by comparing the temperature effect found in evolved genotypes with that found in their ancestral strains. Specifically, we subtracted the expression change in the ancestor from the change observed in EC and EI genotypes (**Figure 2C**). This metric is positive when evolved genotypes show enhanced plasticity relative to their ancestors, and negative when they show reduced or dampened plasticity (**Figure 2C** and **S2C**, **Table S3F**). In *S. cerevisiae*, plasticity was dampened after evolution: EC genotypes showed a smaller change in expression between high temperature and 25 °C for five genes (*RPS6A*, *CRF1*, *SSA4*, *TPK1*, *RLM1*, p < 0.05), and EI genotypes showed a smaller change for *SSA4* and *SLT2* (p < 0.05). In contrast, in *S. eubayanus,* the change in expression was enhanced, but not significantly different from the ancestral strain in both genotypes (EI and EC), indicating that major gene expression changes in *S. eubayanus* are constitutive rather than temperature-dependent (elevated baseline expression at both temperatures). Together, these analyses show that increasing temperature evolution drives opposite regulatory responses in the two species: reduced output and plasticity in *S. cerevisiae* versus elevated, constitutive output in *S. eubayanus*. These functional patterns are congruent with the genomic enrichments in TORC1/PKA/MAPK and support a model in which increasing temperature evolution reprograms central signaling differently in warm-vs. cold-tolerant *Saccharomyces* species.

### Copy number variation associated with thermal evolution

We next evaluated copy number variation (CNV) across all 256 evolved genotypes from all eight species to identify structural genomic changes associated with thermal adaptation (**Figure S3** and **Table S4**). After filtering out telomeric regions and retaining only significant events (|log_2_| ≥ 0.5, FDR < 0.05), we detected only a small number of condition-exclusive CNVs across all species (21 and 22 for increasing and constant temperature conditions, respectively, **Table S4C**), indicating that large genomic rearrangements were rare during thermal evolution.

The amplitude of CNV events was generally modest, with median |log_2_| values around 0.7-1.0 in both evolutionary regimes (**Figure S3A**), suggesting moderate copy number shifts rather than whole-chromosome aneuploidies^35,36^. CNV amplitudes did not differ significantly between constant and increasing temperature conditions (Wilcoxon rank-sum test, p = 0.33), and gains were more frequent than losses across most species in both conditions (**Figure S3B**).

The total number of CNV events varied among species and temperature regimes (**Figure S3C**). Under increasing temperature conditions, CNVs were scattered across multiple chromosomes, with a modest recurrence of events in chromosome XV in *S. kudriavzevii*, *S. mikatae*, and *S. cerevisiae* (**Table S4C**). Under constant temperature, CNVs were predominantly located on chromosomes X and XII, with both genetically different *S. eubayanus* strains sharing a CNV on chromosome X, suggesting a species-specific recurrence. Interestingly, chromosome IX contained CNVs in both temperature conditions but in different species (**Table S4C**), pointing at independent structural changes rather than convergent evolutionary events.

Overall, CNVs were generally rare and species-specific, with no evidence of recurrent, convergent events across species, suggesting that thermal evolution in *Saccharomyces* is primarily driven by point mutations and regulatory changes rather than by large-scale copy-number variation, within the timeframe of this evolution experiment.

### Loss of mitochondrial genome as an adaptive strategy under thermal stress

We detected a consistent loss of mitochondrial DNA (mtDNA) in genotypes evolved under increasing temperature conditions (**Figure 3A** and **S4A**, **Table S4D**). Clear differences emerged between species with different thermal tolerances: all three cold-tolerant species (*S. eubayanus*, *S. kudriavzevii*, and *S. arboricola*) consistently showed a complete loss of mtDNA under increasing temperature conditions in all 31 genotypes we sequenced. In contrast, species with intermediate thermal tolerance (*S. uvarum* and *S. jurei*) showed mixed results, with some genotypes retaining and others losing their mtDNA (all genotypes of the strains Su-1 and Sj-1 lost mtDNA, only 72% of Sj-2 genotypes lost mtDNA, and all Su-2 genotypes retained mtDNA (**Figure 3A** and **S4A**, **Table S4E**). Warm-tolerant species (*S. mikatae*, *S. paradoxus*, and *S. cerevisiae*) also showed a mixed pattern, with all Sm-2 and Sp-2 genotypes completely losing their mtDNA, and 88% Sp-1, 15% Sc-1, and 57% Sc-3 genotypes losing their mtDNA. On the other hand, all Sm-1 and Sc-2 genotypes retained their mtDNA. Notably, under constant-temperature conditions, mitochondrial loss was rare, observed only in the Sk-1 strain and in a single Sp-2 genotype. Together, these results reveal a strong association between thermal regime, species-specific thermal tolerance, and mtDNA stability, with cold-tolerant species showing a markedly higher propensity for mtDNA loss when adapting to increasing temperature.

**Figure 3.**
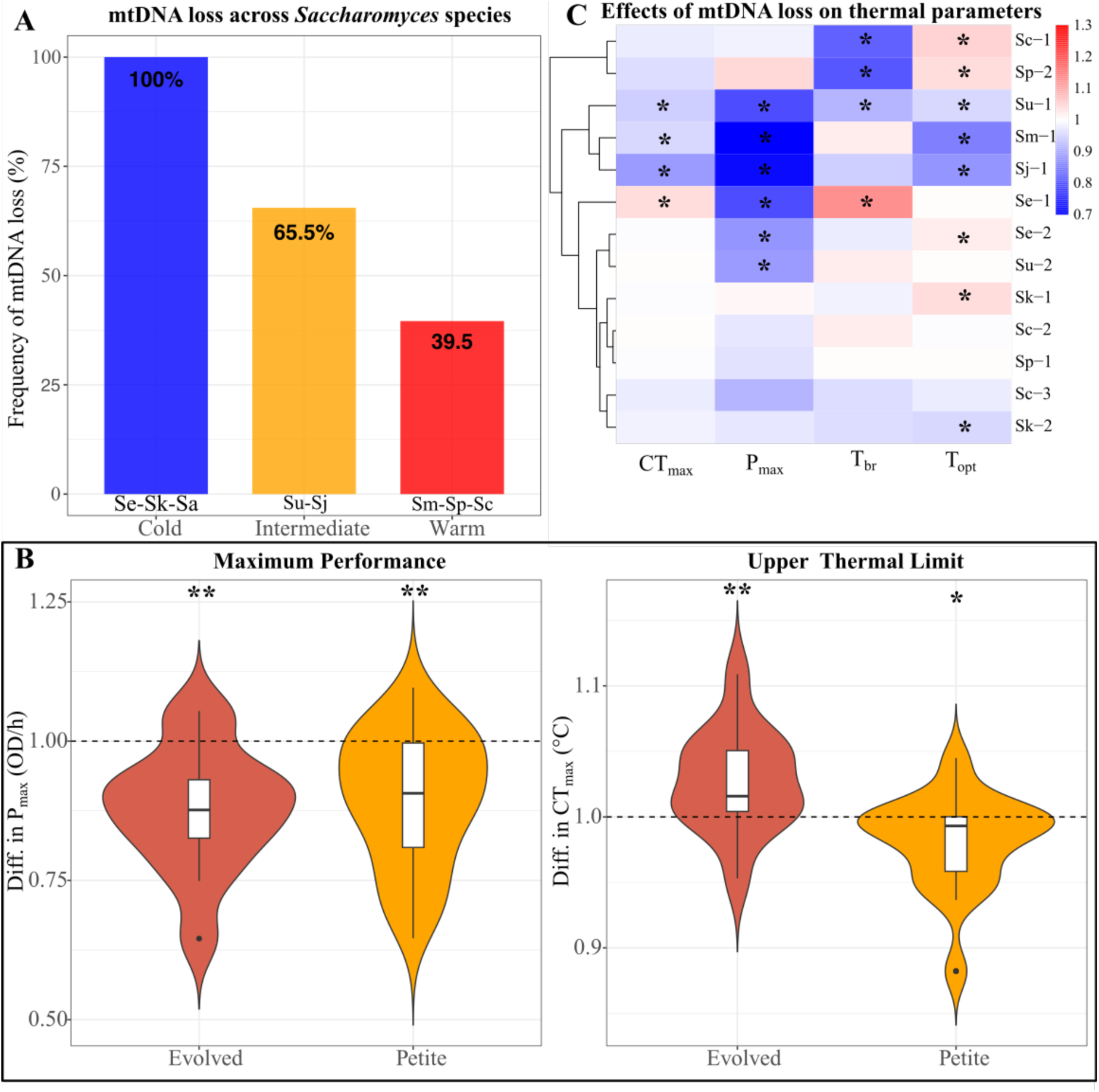
The effect of mtDNA loss on thermal performance across *Saccharomyces* species. **(A)** Frequency of mitochondrial genome loss (petite formation) across *Saccharomyces* species grouped by thermal tolerance, according to Molinet & Stelkens (2024). Cold-tolerant species: *S. eubayanus*, *S. arboricola*, *S. kudriavzevii*; Intermediate species: *S. uvarum* and *S. jurei*; warm-tolerant species: *S. mikatae*, *S. paradoxus*, *S. cerevisiae*. **(B)** Differences in maximum performance (P_max_) and upper thermal limit (CT_max_) between genotypes (Evolved vs Ancestral, Petite vs Ancestral). Violin plots show the distribution of the difference in P_max_ and CT_max_ across strains, with internal boxplots indicating medians and interquartile ranges. The dashed line indicates the performance of ancestral strains, set at 1. Asterisks indicate significant differences between evolved and petite strains relative to their ancestors (Wilcoxon test; * p < 0.05, ** p < 0.001). **(C)** Heatmap showing the relative effects of mtDNA loss on thermal parameters across strains. Values represent the ratio between petite and ancestral strains for each parameter (petite/ancestral). Values < 1 indicate a decrease and values > 1 indicate an increase relative to the ancestors. Hierarchical clustering was applied to strains based on Euclidean distance of scaled values to highlight similarities in response patterns. Asterisks denote significant differences between petite and ancestral strains based on non-overlapping confidence intervals. Color scale indicates the magnitude and direction of change.

To evaluate the role of mitochondria in thermal tolerance and to test whether mtDNA loss or gain alters the shape of thermal performance curves (TPCs), we generated mitochondrial-deficient (petite) strains from each ancestor and compared their growth performance (maximum growth rate, μ_max_) with ancestral and evolved genotypes. All strain types (ancestral, evolved, and petite) were grown at 11 temperatures ranging from 16 to 40 °C (**Table S5A**). We then fitted TPCs for each strain type (ancestral, evolved, and petite) within each strain, plotting the μ_max_ against temperature (**Figure S5**). We first tested whether growth performance differed among strain types (ancestral, evolved, and petite) across the whole temperature range using a linear-mixed-effects model that included temperature as a spline term to capture the shape of the TPC. The model revealed a strong effect of temperature on growth rate (F_3,2055_ =339.0, p < 0.001) and a significant main effect of strain type (F_2,2055_ = 6.30, p = 0.0019), indicating overall differences in thermal performance among ancestral, evolved, and petite strains. The strain type x temperature interaction was marginal (F_6,2055_ = 1.06, p = 0.068), suggesting similar curve shapes (**Table S5B**). Pairwise post hoc comparisons showed that these differences were most pronounced at higher temperatures (>25 °C; **Table S5C**).

Then, we applied a linear model to each strain separately to determine whether the observed differences were general across the genus or strain-specific. Significant effects of strain type were detected in several cold- and intermediate-tolerant strains (e.g. *S. eubayanus* Se-1, *S. uvarum* Su-1, *S. kudriavzevii* Sk-2, and *S. jurei* Sj-1), indicating altered growth performance in mtDNA-depleted strains (**Table S5D** and **S5E**). However, other cold- and intermediate-tolerant strains (*S. eubayanus* Se-2, *S. kudriavzevii* Sk-1, and *S. uvarum* Su-2), together with all strains of the warm-tolerant species (*S. cerevisiae* and *S. paradoxus*), showed no significant performance differences between ancestral and petit strains, suggesting a limited impact of mitochondrial function on thermal performance (**Table S5D** and **S5E**). These results suggest that mitochondrial function affects the shape of TPCs, but that this is strongly strain-dependent.

Analyzing individual thermal performance parameters (T_opt_: optimum temperature, CT_max_: critical thermal maximum, CT_min_: critical thermal minimum, P_max_: maximum performance, T_br_: thermal breadth) across all strains and species showed a significant decrease in petite strains of 10.4% in P_max_ and 1.8% in CT_max_, relative to their ancestors (Wilcox test, p-value = 5.00×10^-3^ and 0.018, respectively, **Figures 3B** and **S4B**, **Tables S5F** and **S5G**), but no significant differences in T_opt_, CT_min_, or T_br_. This suggests that, in general, mtDNA loss constrains both maximal performance and upper thermal limits. This dissociation indicates that the adaptive thermal phenotypes of evolved genotypes require additional regulatory and metabolic reprogramming beyond the loss of respiratory function.

However, global comparisons can mask strain-specific responses. We therefore evaluated the effects of mtDNA loss on thermal parameters also at the strain level (**Figure 3C**, **Table S5H**). Here, the effects of mtDNA loss were highly strain-dependent, revealing four distinct response groups. First, three warm-tolerant petite strains (Sc-2, Sc-3, Sp-1) showed no changes in any thermal parameters relative to their ancestors, indicating that mitochondrial function does not measurably influence TPC shape or thermal limits in these genetic backgrounds. Second, one warm-tolerant and two intermediate-tolerant petite strains (Sm-1, Su-1, and Sj-1, respectively) displayed a consistent pattern of reduced performance and reduced thermal tolerance, characterized by simultaneous decreases in P_max_, CT_max_, and T_opt_. This pattern indicates that, in these backgrounds, mtDNA loss not only lowers maximal growth performance but also shifts the TPC toward lower temperatures and reduces heat tolerance. Third, Se-1 represented a distinct case in which mtDNA loss in the ancestral strain was associated with a partial shift toward higher thermal tolerance despite a cost in performance: P_max_ decreased, while both CT_max_ and T_br_ increased. In this strain, mtDNA loss resulted in an expansion of the thermal range and an elevated upper thermal limit, accompanied by reduced maximal performance (**Figure S5**). Thus, Se-1 petite ancestral strains showed a phenotype similar to that of evolved genotypes, i.e., consistent with the generalist or “a jack of all temperatures is a master of none” pattern^5^. Fourth, mtDNA loss in Sc-1 and Sp-2 was characterized by an increase in T_opt_ coupled with a decrease in T_br_, consistent with a more specialized TPC (a higher optimum but narrower thermal breadth) rather than a broader tolerance to high temperatures.

Together, these results demonstrate that mitochondrial function plays an important, but highly strain-dependent role in shaping thermal performance. While the loss of mtDNA consistently reduced overall performance (P_max_), its impact on thermal optima, upper thermal limits, and thermal breadth varied substantially across strains. Thus, mtDNA loss alone does not explain the evolutionary shifts in thermal tolerance across all species and strains.

## DISCUSSION

Temperature is a key factor influencing cellular physiology, ecological niches, and species distributions^12^. Using a comparative framework across eight *Saccharomyces* species spanning the genus’s full thermal niche, we show that thermal adaptation to sustained temperature increase is characterized by a dual pattern: convergent selection on conserved molecular targets coupled with divergent regulatory and physiological outcomes. By integrating experimental evolution, genome-wide analyses, functional validation, mitochondrial manipulations, and thermal performance assays, our study reveals how climate warming can drive predictable molecular responses while simultaneously amplifying natural species- and strain-specific constraints.

### Convergent molecular targets under thermal selection

Our experimental evolution regime under increasing temperature repeatedly targeted conserved signaling pathways regulating growth, metabolism, and stress responses, particularly TORC1, PKA, and MAPK (**Figure 4A**). Although evidence for parallel evolution at the molecular level in nature is relatively rare (but see, e.g., ^37–39)^, microbial experimental evolution studies often report the independent emergence of mutations in the same gene, protein complex, or regulatory pathway^11,25,40–44^. However, this pattern has largely been limited to experiments initiated from clonal populations. Here, we show that this convergence extends across species with contrasting thermal niches: across the *Saccharomyces* genus, thermal selection consistently targets central regulatory hubs rather than primarily affecting temperature-specific enzymes. This reveals cross-species predictability at the level of conserved regulatory pathways without strict gene-level identity, highlighting shared molecular constraints on thermal adaptation across the *Saccharomyces* genus. The repeated targeting of the same signaling hubs across species, despite distinct genomic backgrounds, suggests non-random selection on conserved regulatory architectures rather than mutation-rate-driven coincidence^16,45^.

**Figure 4.**
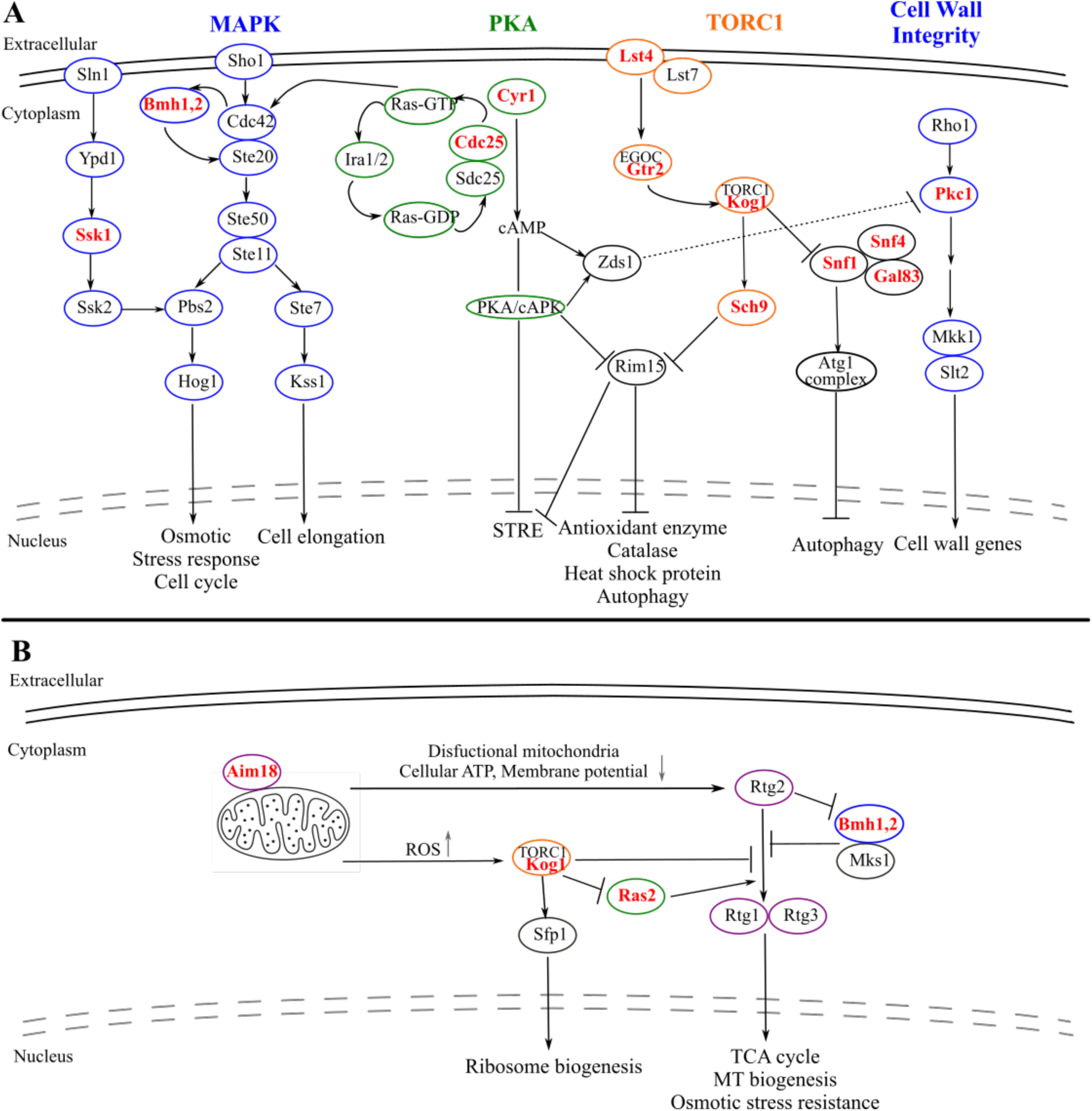
Conserved signaling pathways and mitochondrial feedback shape divergent thermal adaptation trajectories in *Saccharomyces*. **(A)** Schematic representation of conserved growth-stress signaling pathways recurrently targeted during experimental evolution under increasing temperatures. Independent lineages of *Saccharomyces* species accumulated mutations in key components of the TORC1, PKA, and MAPK signaling networks, suggesting these pathways are central regulatory hubs under thermal selection. Only major nodes relevant to repeatedly mutated genes and pathway integration are shown. These conserved pathways coordinate cellular processes, including growth, stress responses (STRE), autophagy, ribosome biogenesis, and cell wall integrity, illustrating how thermal evolution repeatedly targets shared molecular architectures rather than temperature-specific enzymes. **(B)** Conceptual model illustrating the interaction between mitochondrial function and conserved signaling pathways during thermal adaptation. Mitochondrial dysfunction, including mitochondrial genome loss, alters cellular energy status, membrane potential, and redox balance, generating signals (e.g., changes in ATP levels and reactive oxygen species) that feed back into central regulatory circuits such as TORC1 and RAS-PKA signaling. Retrograde signaling components (e.g., Rtg1/3 and Rtg2) further link mitochondrial status to nuclear gene expression. These interactions modulate, but do not solely determine, adaptive outcomes, contributing to strain- and species-specific differences in thermal performance. Genes with mutations identified in evolved genotypes are shown in red. The mitochondrial image in B was generated with AI. Adapted from ^16,33,46^.

Recurrent mutations in key regulatory genes such as *CDC25*, *CYR1*, *BMH1*, and *SNF4* highlight the importance of nodes that integrate nutrient sensing, growth, and stress signaling. These genes occupy central positions within their respective pathways and are well-suited to mediate trade-offs between proliferation and stress resistance^33,46–48^. Importantly, many strain-specific mutations not shared across species mapped to the same signaling networks, suggesting that thermal adaptation can proceed through multiple genetic routes that converge at the level of pathway output. Together, these patterns support a model in which thermal evolution is constrained by the architecture of conserved signaling systems, favoring repeated modification of the same regulatory modules across independent lineages.

### Divergent regulatory outcomes despite molecular convergence

While genomic analyses revealed strong convergent evolutionary patterns in TORC1, PKA, and MAPK pathways, our functional validation demonstrated that this convergence does not translate into uniform regulatory outcomes. Instead, when assayed near the thermal limits of warm- and cold-tolerant species, evolution under increasing temperature reprogrammed pathway activity in opposite directions, depending on the species background. Ancestral genotypes (i.e., before experimental evolution) of warm- and cold-tolerant species already differed markedly in baseline signaling output and in their transcriptional responses to heat stress, reflecting distinct regulatory architectures shaped by their thermal adaptation.

These differences in pathway activity were further accentuated by experimental evolution. In the warm-tolerant species *S. cerevisiae*, genotypes evolved under increasing temperature showed reduced pathway output and dampened transcriptional plasticity at high temperature, consistent with regulatory buffering of an already heat-adapted organism. In contrast, in the cold-tolerant species *S. eubayanus*, evolved genotypes showed strong, often constitutive upregulation of the same pathway targets, indicating a shift toward sustained stress signaling rather than plastic regulation. Thus, although thermal selection repeatedly targets the same signaling hubs, adaptive solutions exploit different regions of the regulatory landscape depending on species-specific constraints. This functional divergence despite shared molecular targets underscores the importance of genetic background in shaping adaptive trajectories. Rather than converging on a single optimal signaling state, thermal adaptation appears to involve context-dependent rewiring of conserved pathways, reflecting different balances between growth and stress resistance.

### Mitochondrial function as a context-dependent constraint for thermal adaptation

A second major axis of divergence in our study involves mitochondrial function. Loss of mtDNA occurred frequently in populations evolved under increasing temperature and was universal in cold-tolerant species. Similar patterns of mtDNA instability or loss under thermal stress have been reported in previous experimental evolution studies in yeast, suggesting that thermal stress can destabilize mitochondrial maintenance or select against respiratory metabolism^16,19^. Mitochondria play a central role in energy production, redox balance, and stress signaling^49^, and high temperatures are known to challenge mitochondrial integrity by increasing membrane fluidity, disrupting electron transport, and elevating reactive oxygen species^50–52^. Consistent with this, multiple studies have shown that mitochondrial genotype and function strongly modulate thermal tolerance^19,29,53–56^. In cold-tolerant species, mitochondrial dysfunction may represent a recurrent, possibly adaptive, response to increasing temperature stress. Loss of respiration may reduce oxidative stress or mitigate metabolic imbalance at elevated temperatures, as has been proposed in other contexts of thermal or oxidative stress^53^.

Despite prevailing mtDNA loss, our functional assays demonstrate that mitochondrial deficiency alone is insufficient to cause significant fitness gains or adaptive shifts in thermal tolerance. Across species, loss of mtDNA in ancestral strains consistently reduced maximal growth performance, indicating a substantial energetic cost. Moreover, our analyses revealed that petite strains exhibited, on average, a reduction in Ct_max_, rather than an increase, relative to their ancestral counterparts, arguing against a simple or universally adaptive role for mtDNA loss during thermal evolution. These results are in line with recent work showing that mitochondrial perturbations can constrain thermal tolerance by limiting bioenergetic capacity or disrupting mitochondrial-nuclear coordination^57^.

Importantly, effects of mtDNA loss on thermal parameters were highly strain-dependent: Only a single cold-tolerant petite strain (Se-1) showed partial phenotypic convergence toward the heat tolerance of the genotypes evolved in increasing temperature, resembling the characteristic generalist profile or “a jack of all temperatures is a master of none” phenotype. A subset of warm-tolerant strains (Sc-2, Sc-3, and Sp-1) showed no detectable changes in any thermal parameter following mtDNA loss, while other warm-tolerant strains (Sc-1 and Sp-2) showed a thermal specialization rather than a “hotter is wider” phenotype.

These results indicate that mitochondria act as modulators rather than primary drivers of thermal adaptation. Although mtDNA loss is a recurrent outcome of evolution under increasing temperature in yeast^16,19^, its phenotypic consequences are highly context-dependent, ranging from neutral to deleterious, and only rarely mimicking the adaptive shifts observed after experimental evolution. Differences in mitochondrial–nuclear coordination, redox homeostasis, or retrograde signaling likely determine whether mitochondrial dysfunction alleviates or exacerbates thermal stress^58^. Notably, several of the conserved signaling pathways repeatedly targeted by thermal selection are known to sense mitochondrial status and metabolic flux^46^, suggesting that mitochondrial function feeds back into the same regulatory circuits that shape adaptive thermal responses.

### Implications for predicting evolutionary responses to climate warming

Together, our findings support a multi-layered model of thermal adaptation in which climate warming imposes shared molecular constraints while generating divergent physiological outcomes. Increasing temperature consistently selects for changes in conserved signaling hubs, but the adaptive consequences of these changes depend on species-specific regulatory architectures and energetic constraints. As a result, similar molecular signatures can give rise to distinct thermal performance curves and adaptive strategies, even within a single genus.

From an ecological perspective, our results highlight the challenges of predicting evolutionary responses to climate warming. Adaptive potential cannot be inferred from molecular changes alone, nor from species-level classifications, but instead emerges from the interaction between conserved stress-response pathways, organelle function, and genetic background. The frequent loss of mtDNA under thermal stress further underscores the role of energetic trade-offs in shaping evolutionary trajectories, particularly in environments with recurrent high temperatures.

More broadly, our study suggests that climate change is likely to promote both convergent molecular evolution and divergent ecological outcomes, reshaping microbial diversity through a mosaic of adaptive solutions. Integrating genomic, functional, and physiological approaches will therefore be essential for understanding and predicting how organisms respond to ongoing global warming.

## MATERIALS AND METHODS

### Strains

We used strains from an experimental evolution study that exposed populations of eight *Saccharomyces* species (*S. cerevisiae*, *S. paradoxus*, *S. mikatae*, *S. jurei*, *S. kudriavzevii*, *S. arboricola*, *S. uvarum*, and *S. eubayanus*) to gradually increasing temperature (25-40 °C) and constant temperature (25 °C) for up to 600 generations^5^. We selected two genotypes from each replicate evolution line per strain and temperature condition. We used four replicate lines per strain from a total of 16 strains (2 per species), and the two temperature conditions, yielding a total of 256 genotypes. These genotypes, together with their respective ancestral strains, were sequenced for whole-genome analysis (**Table S1**). Ancestral strains represent the starting point before experimental evolution (16 strains), evolved genotypes under constant temperature (EC) were used as thermal control (128 genotypes), and evolved genotypes under increasing temperature (EI) represent heat-adapted lineages (128 genotypes). All strains were maintained on YPD agar (1% yeast extract, 2% peptone, 2% glucose, and 2% agar) and stored at −70 °C in 20% glycerol stocks.

### Whole genome analysis

Genomic DNA was obtained for whole-genome sequencing using the Thermo Scientific KingFisher^TM^ Duo Prime Purification system, and sequenced at the Earlham Genomics facilities (https://www.earlham.ac.uk/) using the LITE pipeline^59^. The quality of the raw sequencing reads was assessed using FastQC 0.12.1 (https://www.bioinformatics.babraham.ac.uk/projects/fastqc/) and MultiQC 1.22.2^60^, both before and after trimming and quality filtering using fastp 0.23.4^61^. Reads were aligned against reference genomes of each species (**Table S2A**) using BWA 0.7.18^62^. BAM files were sorted using SAMtools 1.20^63^. Duplicate reads were removed using Picard 2.27.5 (https://broadinstitute.github.io/picard/), and mapping quality was assessed using QualiMap 2.2.1^64^ (**Table S2B**).

### SNPs and CNV analysis

Variant calling and filtering were performed using GATK version 4.3.0.0^65^. Variants were called per sample using HaplotypeCaller (default settings), generating g.vcf files. Variant databases were constructed using GenomicsDBImport, and genotypes were called using GenotypeGVCFs with the -G Standard Annotation option. SNPs and InDels were extracted and filtered out separately using SelectVariants. We then applied recommended filters with the following options: QD < 2.0, FS > 60.0, MQ < 40.0, SOR > 4.0, MQRankSum < −12.5, ReadPosRankSum < −8.0. This vcf file was further filtered by removing missing data using the option-max-missing 1, filtering out sites with a coverage below the 5^th^ or above the 95^th^ coverage sample percentile using the options-min-meanDP and -max-meanDP, and a minimum site quality of 30 (-minQ 30) in VCFtools 0.1.16^66^. Sites with a mappability score less than 1, as calculated by GenMap 1.2.0^67^, were filtered using bedtools 2.18^68^. As an additional filtering step, the ancestral and evolved files were intersected using BCFtools 1.20^69^, and variants with shared positions were extracted from the vcf files of the evolved genotypes. Annotation and effect prediction of the variants were performed with SnpEff^70^. To enable cross-species comparison, gene identities were assigned based on species-specific genome annotations when available, or inferred by BLAST-based homology against the *S. cerevisiae* reference genome. Gene ontology analysis was performed using the tools provided by the DAVID Bioinformatics Resource^71^, selecting categories with a significant overrepresentation using an FDR < 0.05.

CNVs were called using CNVkit (whole-genome mode)^36,72^ from the Illumina WGS data of all ancestral and evolved genotypes within each species/strain background. To ensure identical binning across samples, including the mitochondrial chromosome, we first created an accessibility BED file from the species reference (FASTA files). Next, we generated a single whole-genome bin set per species using 1-kb tiles and used the same BED file and reference for every sample of that species. For each BAM file, we computed coverage in the 1-kb bins, normalized to the species reference and ancestral strain, and segmented into contiguous genomic regions with similar normalized log_2_ copy-ratio profiles. For each sample, we computed per-segment confidence intervals and t-tests, and merged these metrics back with the segments. Segments were kept as significant if all of the following held: the 95% CI on log_2_ fold-change did not cross zero, Benjamini-Hochberg FDR on the t-test < 0.05 (within sample), did not correspond to telomeric regions, and the effect size was biologically meaningful for a diploid genome (amplitude thresholds corresponding to ± 1 copy: gain if log_2_ ≥ +0.50, loss if log_2_ ≤-1.00; +2 copies, when log_2_ ≥ +1.00). For gains and losses separately, we reduced all significant segments into consensus intervals and computed a sample-by-region presence matrix, calling a region “present” in a sample if ≥ 50% of the consensus interval overlapped that sample’s significant segment. Regions were then classified as exclusive to the evolved under increasing temperature genotypes (present in the ≥1 evolved genotype, absent in the evolved genotypes under constant temperature) or exclusive to the evolved under constant genotypes (present in the ≥1 evolved genotype, absent in the evolved genotypes under increasing temperature).

To quantify mtDNA copy change per sample, we used the corrected bin-level files (.cnr). After excluding bins with zero depth and extreme outliers, we computed the median log_2_ ratio across mtDNA bins and subtracted the median across nuclear DNA bins from the same sample, yielding a relative mtDNA log_2_ (mt – nuclear). Samples were heuristically classified as: mtDNA loss when the relative mtDNA log_2_ was ≤ −1.0 (≥ 2x lower than nuclear), mtDNA depletion when the relative mtDNA log_2_ was between −1.0 and −0.6, mtDNA normal-like when the relative mtDNA log_2_ was between −0.6 and +0.4, and mtDNA enrichment candidate when the relative mtDNA log_2_ was ≥ +0.4. mtDNA loss was visually verified in the Integrative Genomics Viewer (IGV) software, and by growing the genotypes on YPG agar (1% yeast extract, 2% peptone, 2% glycerol, 2% agar), where growth requires a functional mitochondrial respiratory chain and thus fails in mitochondrial genome-deficient (petite) genotypes.

### Generation of rho^0^ strains

Strains lacking mtDNA (rho^0^) were generated by treating ancestral strains (rho^+^) with ethidium bromide^73^. Briefly, cells were grown overnight in YPD at 25 °C. Then, 100 μL of the culture was diluted in 1 mL of YPD and incubated at 25 °C until it reached the exponential growth phase (OD_600nm_ 0.4-0.6). Cells were washed and resuspended in 0.1 M potassium phosphate buffer at a concentration of 10^6^ cells/mL. Ethidium bromide was added to a final concentration of 10 μg/mL and incubated for 8 h. After incubation, 200 μL of the cell culture was plated on YPD agar and incubated at 25 °C for 3-4 days. The mtDNA loss was evaluated by growing the colonies in YPG agar and by standard colony PCR using primers previously described for mtDNA sequences^29^ (**Table S3**). Because of the mutagenic nature of ethidium bromide, we controlled for the potential effect of spurious nuclear mutations by using three rho^0^ colonies (replicates) from each ancestral strain (**Table S1D**). We did not obtain rho^0^ colonies for *S. mikatae* yHAB336 (Sm-2), *S. jurei* NCYC3947 (Sj-2), and *S. arboricola* ZP960 (Sa-1), despite repeated attempts using the ethidium bromide protocol, suggesting strain-specific resistance to mitochondrial genome loss under these conditions.

### Thermal performance curves analysis

Strains were phenotypically characterized under microculture conditions as previously described^5^. Mitotic growth was measured in 96-well plates at 11 different temperatures: 16, 18, 20, 23, 25, 28, 31, 32, 34, 37, and 40 °C. Inocula were prepared by growing one colony of each strain for 24 h at 25 °C in 96-well plates with 200 μL YPD in each well. Cultures were then diluted to an initial OD_600nm_ of 0.1 in fresh YPD for growth at the 11 temperatures for 24-48 h until all strains had entered the stationary phase. The next day, these cultures were used to inoculate a new 96-well plate with 200 μL of YPD per well, at an initial OD_600 nm_ of 0.1, and kinetic growth parameters were measured. Growth curves were obtained by measuring OD_600nm_ every 30 min in a TECAN Sunrise instrument at all 11 temperatures for 24-48 h, until all strains had entered the stationary phase. Three independent OD measurements were taken per strain (i.e. technical triplicates). Maximum growth rate (μ_max_) was determined as previously described^74^. For this, μ_max_ was calculated following a smoothing procedure on ln-transformed OD-values and using the discrete derivative, as previously described^75^ in R version 4.3.1.

We fitted thermal performance curves to μ_max_ of each strain using the cardinal temperature model with inflection (CTMI) as previously described^6^, using equation 1.

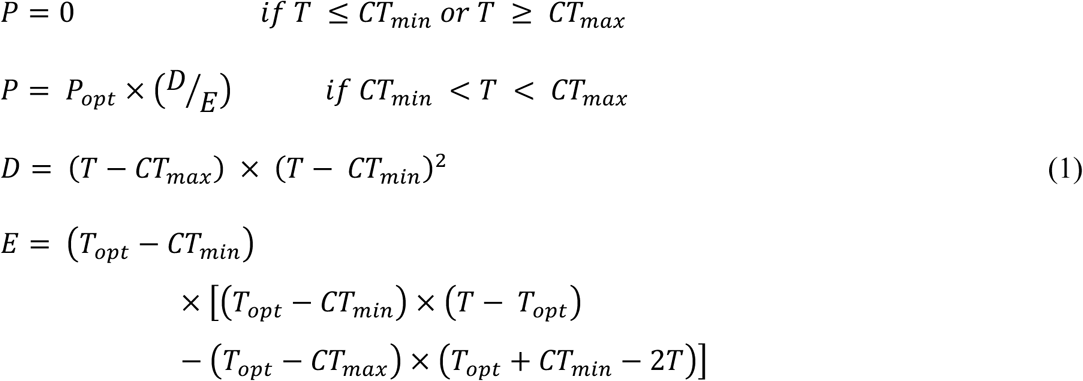

Where:

CT_max_ is the temperature above which no growth occurs. CT_min_ is the temperature below which no growth occurs.

T_opt_ is the temperature at which P_max_ equals its optimal value (P_opt_).

CTMI parameters were estimated using nonlinear regression in R version 4.3.1. Thermal tolerance and T_br_ were obtained using the calc_params function from the rTPC package^76^, where thermal tolerance corresponds to CT_max_ minus CT_min,_ and thermal breadth (T_br_) is the temperature range across which performance is above 80% of optimal. The adequacy of fit of TPCs was checked by the proportion of variance explained by the model (R^2^) and by the residual sum of squares (RSS). Relative thermal parameters were obtained by normalizing each parameter in the petite and evolved strains to their corresponding ancestral values and used to construct a heatmap with hierarchical clustering. Hierarchical clustering was performed on strains using Euclidean distance and complete linkage. Heatmap visualization was generated using the pheatmap package. We considered thermal parameters to differ significantly among strains if their 95% confidence intervals did not overlap.

### RNA extraction and qPCR assay

Gene expression of target genes in the TORC1 (*RPS6A*, *CRF1*), PKA (*SSA4*, *TPK1*), and MAPK (*SLT2*, *RLM1*) signaling pathways was evaluated for ancestral strains, and genotypes evolved under increasing and constant temperatures in the species *S. cerevisiae* and *S. eubayanus* (strains Y12 (Sc-1) and CBS12357 (Se-1), respectively). Gene expression analysis was performed by qPCR from exponential cultures grown at 25 °C for both species, 34 °C for *S. eubayanus* strains, and 40 °C for *S. cerevisiae* strains. These temperatures correspond to the maximum reached during the experimental evolution for each strain. This design intentionally probed each species near its upper thermal limit, allowing comparison of regulatory responses under ecologically relevant heat stress. Cells were first grown in 5 mL of YPD for 24 h at 25 °C and then diluted to an initial OD_600 nm_ of 0.1 in fresh YPD for growth at the three different temperatures of interest, until they reached the exponential phase (OD_600 nm_ of ∼0.6). Cells were collected, immediately frozen in liquid nitrogen, and stored at −70 °C until RNA extraction. RNA was extracted using the YeaStar^TM^ RNA Kit (Zymo Research) according to the manufacturer’s instructions. Then, genomic DNA traces were removed by treating samples with DNase I (Zymo Research), and total RNA was recovered using the RNA Clean and Concentrator^TM^ kit (Zymo Research). Concentrations of the purified RNA were determined using a UV-Vis spectrophotometer and verified on 1.5% agarose gels. The RNA extractions were performed in three biological replicates.

cDNA was synthesized using 200 units of RevertAid M-MulV RT (Thermo Scientific), 0.5 μg of Oligo (dT)_18_ primer, and 0.5 μg of RNA in a final volume of 20 μL according to the manufacturer’s instructions. The qPCR reactions were carried out using Maxima SYBR Green/ROX qPCR Master Mix (Thermo Scientific) in a final volume of 12 μL, containing 0.3 μM of each primer and 1 μL of the cDNA previously synthesized. The qPCR reactions were carried out in two technical replicates per biological replicate using a Step One Plus Real-Time PCR System (Applied Biosystems) under the following conditions: 95 °C for 10 min, followed by 40 cycles at 95 °C for 15 s and 60 °C for 1 min. The genes and primers used are listed in **Table S6**. The relative expression of the target genes was quantified using the mathematical method described by^77^ and normalized with three housekeeping genes (*ACT1*, *UBC6*, and *RPN2*), following established procedures^78,79^. For each sample, noremalization was based on the median Ct value of the two technical replicates. Statistical analysis was performed on the log-transformed, normalized expression per sample^80^.

### Statistical analysis

Statistical analysis and data visualization were performed in R version 4.3.1. To test whether the accumulation of de novo SNPs differed among species and evolutionary regimes, we modeled the number of SNPs detected per evolved genotype using a generalized linear mixed-effects model (GLMM). Because SNP counts are discrete and overdispersed, we fitted a negative binomial model with a log link function. Species, evolutionary condition (constant vs. increasing temperature), and their interaction were included as fixed effects, while strain was included as a random effect to account for non-independence among genotypes derived from the same ancestral strain. Models were fitted using the glmmTMB package. Statistical significance of mixed effects was assessed using Type II WALD χ^2^ tests. When significant interactions were detected, post hoc contrasts comparing evolutionary conditions within each species were performed using estimated marginal means (emmeans), with p-values adjusted for multiple testing using the Benjamini-Hochberg false discovery rate (FDR). Model assumptions were evaluated by inspecting residual distributions and dispersion parameters.

For qPCR analyses and to keep comparisons statistically coherent, we used two complementary normalizations: for comparison between ancestral *S. cerevisiae* and *S. eubayanus* at 25 °C, all samples were calibrated to the *S. cerevisiae* ancestral strain at 25 °C; for all within-species contrasts (25 °C vs high temperature; ancestral vs evolved at a given temperature), each species was calibrated to its own ancestral strain at 25 °C or high temperature condition (40 °C for *S. cerevisiae* and 34 °C for *S. eubayanus*). All gene-level tests were run per gene and (when applicable) per species using Welch’s t-test on log_2_ normalized expression values and corrected using Benjamini-Hochberg false discovery rate (FDR). To test whether the change from 25 °C to high temperature differs between *S. cerevisiae* and *S. eubayanus*, ordinary least squares (OLS) models with robust (HC3) standard errors were fit as log_2_ normalized expression ∼Species * Temperature. To test the change in temperature sensitivity in evolved genotypes, we first computed the temperature effect as expression at high temperature minus expression at 25 °C for each strain. Then, we tested whether evolution altered this temperature sensitivity by fitting per-species OLS models with HC3 as log_2_ normalized expression ∼Species * Evolved genotype. An adjusted p-value < 0.05 was considered significant.

To evaluate whether mtDNA loss affected thermal performance, we analysed μ_max_ for ancestral, evolved, and petit strains across 11 assay temperatures (16–40 °C). First, we tested overall differences in thermal performance across all strains using a linear mixed-effects model fitted with the nlme package (version 3.1-168) as μ_max_ = Genotype x ns(Temperature, df = 3) + (1|Strain), where Genotype had three levels (Ancestral, Evolved, Petite), and ns(Temperature, df = 3) denotes a natural spline function used to capture the non-linear effect of temperature. Models were fitted by restricted maximum likelihood (REML), and heteroscedasticity in the fitted values was modelled using a power variance structure (varPower(form = ∼fitted(.))). The significance of fixed effects (Genotype, Temperature, and their interaction) was assessed by Type III ANOVA. We then determined whether genotype effects were strain-dependent by fitting separate linear models for each strain: μ_max_ = Genotype × ns(Temperature, df = 3). Pairwise contrasts between genotypes (Ancestral-Evolved, Ancestral-Petite, Evolved-Petite) were computed using the emmeans package (version 1.11.2-8) with Benjamini-Hochberg false discovery rate correction. Model assumptions (normality, independence, and homoscedasticity) were verified with the check_model() function from the performance package (version 0.15.1) and were met in all cases.

To compare thermal parameters among genotypes, we used a non-parametric Kruskal-Wallis test followed by Benjamini-Hochberg-corrected Wilcoxon pairwise comparisons, because residuals did not meet normality assumptions. All parameters were previously normalized to the mean of the ancestral genotype within each strain to allow direct comparison across species.

## Data availability statement

All fastq sequences were deposited in the National Center for Biotechnology Information (NCBI) as a Sequence Read Archive under the BioProject accession number PRJNA1270780. The code used for analyses and plotting is available in our GitHub repository, https://github.com/j-molinet/TPC_genomic_paper. All other data are included in the manuscript and/or in the supporting information.

## ACKNOWLEDGMENTS

We thank Chloé Haberkorn for her help in the laboratory. Computation and data handling were enabled by resources provided by the National Academic Infrastructure for Supercomputing in Sweden (NAISS), partially funded by the Swedish Research Council through grant agreement no. 2022-06725, under Project NAISS 2024/22-917 and 2024/23-411. RK, JM, and CG were supported by the Swedish Research Council (2022-03427) and the Knut and Alice Wallenberg Foundation (2017.0163 and 2024.0216). JM and PV were funded by the Agencia Nacional de Investigación y Desarrollo (ANID) Fondecyt Iniciación grants 11260235 and 11240649, respectively, and by the ANID Iniciativa Científica Milenio program ICN17_022. We acknowledge Fundación Ciencia & Vida for providing infrastructure and laboratory space.

## AUTHOR CONTRIBUTIONS

Conceptualization: J.M., R.S.; Investigation: J.M., R.S.; Methodology: J.M., C.G., P.V.; Formal Analysis: J.M., R.S., P.V.; Resources: R.S.; Visualization: J.M.; Original Draft Preparation: J.M., R.S. All authors have read and agreed to the published version of the manuscript.

## Supplementary Information

**Figure S1. Detailed distribution of de novo SNPs across strains and evolutionary conditions in *Saccharomyces*. (A)** Total number of SNPs per strain, grouped by evolutionary condition (constant vs. increasing temperature). **(B)** Distribution of SNPs across coding and non-coding regions per strain. The upper panel corresponds to strains evolved under increasing temperature, while the lower panel shows strains evolved under constant temperature. **(C)** Parallel evolution across species under constant temperature, displayed as an UpSet plot. Species are grouped by thermal tolerance (blue = cold-tolerant, orange = intermediate, red = warm-tolerant).

**Figure S2. Additional validation of TORC1/PKA/MAPK outputs across species and evolutionary conditions. (A)** Log2 normalized expression for ancestral *S. cerevisiae* (SC-A, red) and *S. eubayanus* (SE-A, blue) across the six genes at 25 °C. Values are relative to SC-A. **(B)** Log2 normalized expression of evolved genotypes at 25 °C relative to their species’ ancestral strains at the same temperature (SC top; SE bottom). EC: genotype evolved at a constant 25 °C; EI: genotype evolved under increasing temperatures. For (A) and (B), the dashed line marks no change. Plotted values correspond to the average of the biological replicates (n=3). Asterisks denote between-species differences in (A) and evolved-ancestral differences in (B) (Welch’s t-test, * p < 0.05, ** p < 0.01, *** p < 0.001, **** p < 0.0001). Genes are grouped by pathway (TORC1: *RPS6A*, *CRF1*; PKA: *SSA4*, *TPK1*; MAPK: *SLT2*, *RLM1*). **(C)** Change in expression between each species’ high temperature and 25 °C for A, EC, and EI (computed as high temperature minus 25 °C; high temperature is 40 °C for SC and 34 °C for SE). The horizontal dashed line marks no change. Asterisks denote differences within genotype, per gene and species (Welch’s t-test, * p < 0.05, ** p < 0.01, *** p < 0.001, high vs 25 °C).

**Figure S3. Copy number variation associated with thermal evolution in *Saccharomyces*. (A)** Distribution of CNV amplitudes (absolute log₂ copy-number change) for all significant CNV events detected across species under constant and increasing temperature evolution regimes. Violin plots show the density of CNV amplitudes, with embedded boxplots indicating median and interquartile range. CNV amplitudes did not differ significantly between evolutionary conditions (Wilcoxon rank-sum test, p = 0.33), indicating comparable magnitudes of copy-number changes across regimes. **(B)** Total number of CNV events per species under constant and increasing temperature conditions. Bars represent the number of condition-exclusive CNVs detected in each species, highlighting substantial species-specific variation but no consistent increase in CNV burden under increasing temperature evolution. **(C)** Proportion of CNV gains and losses per species under constant and increasing temperature conditions. Stacked bars show the relative contribution of copy-number gains and losses within each species and condition, illustrating that gains were generally more frequent than losses across taxa.

**Figure S4. Thermal parameters across ancestral, evolved, and petit strains. (A)** Frequency of mitochondrial genome loss (petite formation) across *Saccharomyces* species. **(B)** Differences in thermal parameters between genotypes (Evolved vs Ancestral, Petite vs Ancestral). Violin plots show the distribution of differences in thermal parameters across strains, with internal boxplots indicating medians and interquartile ranges. The dashed line indicates the ancestral strains set at 1. Asterisks indicate significant differences between evolved and petit strains relative to their ancestors (Wilcoxon test; * p < 0.05, ** p < 0.01, ns not significant).

**Figure S5. TPCs of ancestral, evolved, and petit genotypes per strain.** Changes in the maximum growth rates as a function of temperature for each strain, representing the TPCs. Ancestral TPC is in grey, evolved TPC under increasing temperature is in red, and petite TPCs are in green, orange, and purple. TPCs were constructed using the maximum growth rate of three technical replicates and were fitted using the CTMI.

**Table S1. (A)** Ancestral strains used in this study. **(B)** Evolved genotypes under constant temperature (25 °C) used in this study. **(C)** Evolved genotypes under increasing temperatures used in this study. **(D)** Rho0 strains generated in this study.

**Table S2. (A)** List of reference genomes used in this study. **(B)** Bioinformatics summary statistics. **(C)** Total number of SNPs. **(D)** List of SNPs identified in evolved genotypes under increasing temperatures. **(E)** List of SNPs identified in evolved genotypes under constant temperatures. **(F)** Effects of species and evolutionary regime on the accumulation of de novo SNPs. **(G)** GO and KEGG pathway enrichment analysis for genes with de novo mutations.

**Table S3. (A)** Relative gene expression for target genes of TORC1, PKA, and MAPK pathways in S. cerevisiae and S. eubayanus strains at 25 °C and higher temperatures. **(B)** Baseline cross-species comparison at 25 °C. Log2 normalized expression per gen (ancestral strains), Welch’s tests (two sided), 95% CIs, Hedges’g, and BH-FDR. **(C)** Ancestral temperature effect within species (High – 25 °C) and Species*Temperature interaction. Per species and gene: High – 25 °C mean difference (Welch) with 95% CIs and FDR; OLS (HC3) interaction estimates testing whether SC and SE differ in their temperature response. **(D)** Evolved vs. Ancestral at high temperature. Per species and gene: A vs. EC and A vs. EI at high temperature (SC 40 °C; SE 34 °C): Log_2_ mean differences, 95% CIs, Hedges’g, Welch p-values, BH-FDR. **(E)** Evolved vs. Ancestral at 25 °C. Per species and gene: A vs. EC and A vs. EI at 25 °C. Log_2_ mean differences, 95% CIs, Hedges’g, Welch p-values, BH-FDR. **(F)** Change in temperature responsiveness after evolution (difference-in-differences within species). Per species and gene: estimates of “(High-25 in EC/EI)-(High-25 in A)” alongside OLS (HC3) Temperature x Evolution interaction coefficients, 95% CIs, and BH-FDR (these underpin in Fig. 2C).

**Table S4. (A)** Per sample CNV calls (gain/loss) with CI/FDR, log_2_, and size thresholds. **(B)** Condition-exclusive consensus CNVs (Increasing vs. Control evolved conditions) with recurrence and median log_2_. **(C)** Condition-exclusive consensus CNVs (Increasing vs. Control evolved conditions) with recurrence and median log2, excluding telomeric regions (10 kb). **(D)** Per-sample mitochondrial status. **(E)** CNV summary by evolutionary condition.

**Table S5. (A)** Estimated maximum growth rate for each strain and temperature. **(B)** Linear mixed-effects model results for growth rate as a function of temperature and strain type. **(C)** Pairwise comparison of growth rate between strain types at each temperature. **(D)** Linear mixed-effects model results for growth rate for each strain. **(E)** Pairwise comparison of growth rate between strain types for each strain. **(F)** Estimated thermal parameters of the TPCs for each strain and strain type. **(G)** Kruskal-Wallis rank sum test and Wilcoxon test with Benjamini-Hochberg correction for multiple pairwise comparisons for relative thermal parameters for each strain type. **(H)** Overlapping of confidence intervals of thermal parameters.

**Table S6.** List of primers used in this study.

