## Supplementary material for "Convergent targeting of conserved regulatory networks during thermal evolution across *Saccharomyces*": Figure S1

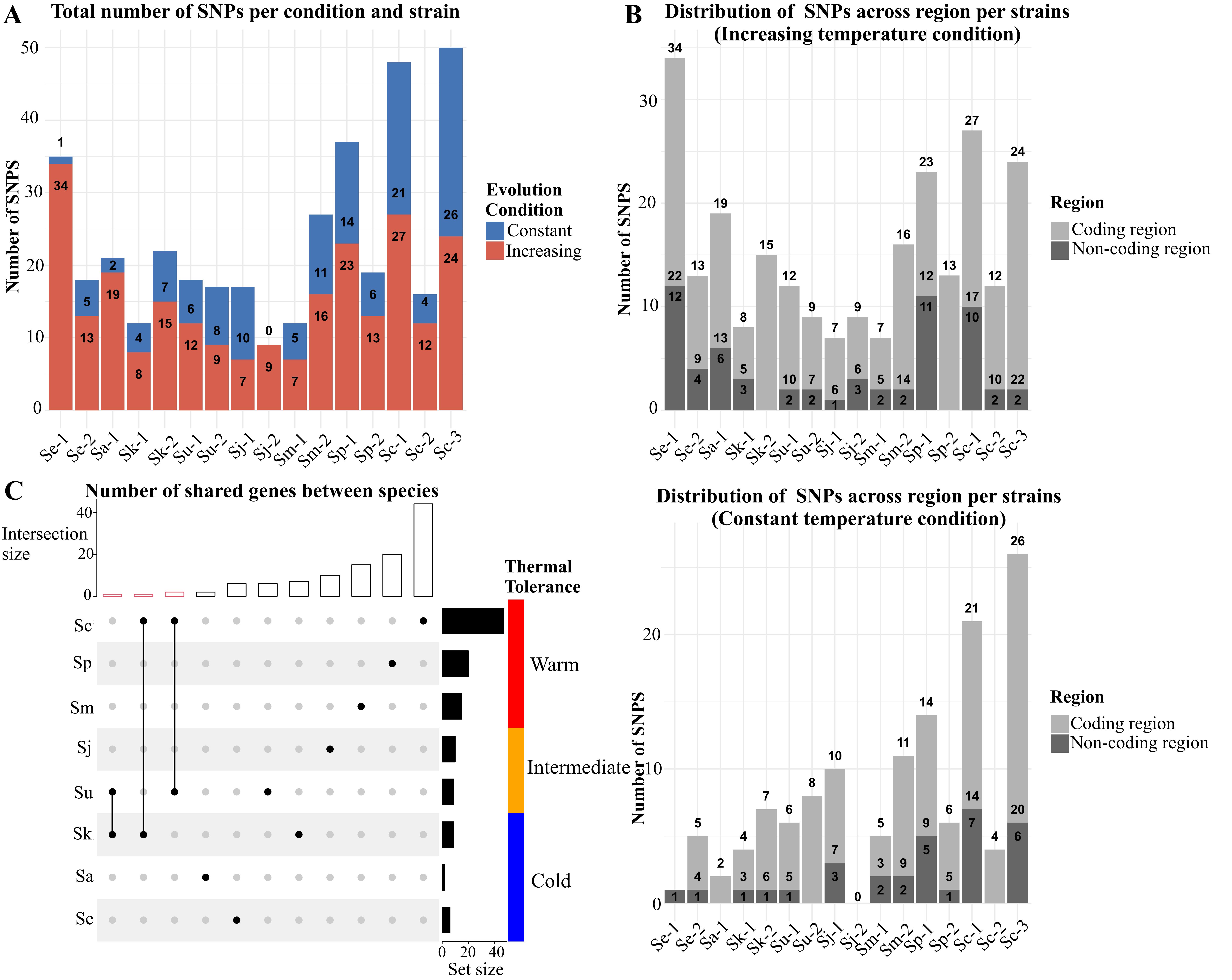


**Figure S1. Detailed distribution of de novo SNPs across strains and evolutionary conditions in *Saccharomyces*. (A)** Total number of SNPs per strain, grouped by evolutionary condition (constant vs. increasing temperature). **(B)** Distribution of SNPs across coding and non-coding regions per strain. The upper panel corresponds to strains evolved under increasing temperature, while the lower panel shows strains evolved under constant temperature. **(C)** Parallel evolution across species under constant temperature, displayed as an UpSet plot. Species are grouped by thermal tolerance (blue = cold-tolerant, orange = intermediate, red = warm-tolerant).
