## Supplementary material for "Convergent targeting of conserved regulatory networks during thermal evolution across *Saccharomyces*": Figure S2

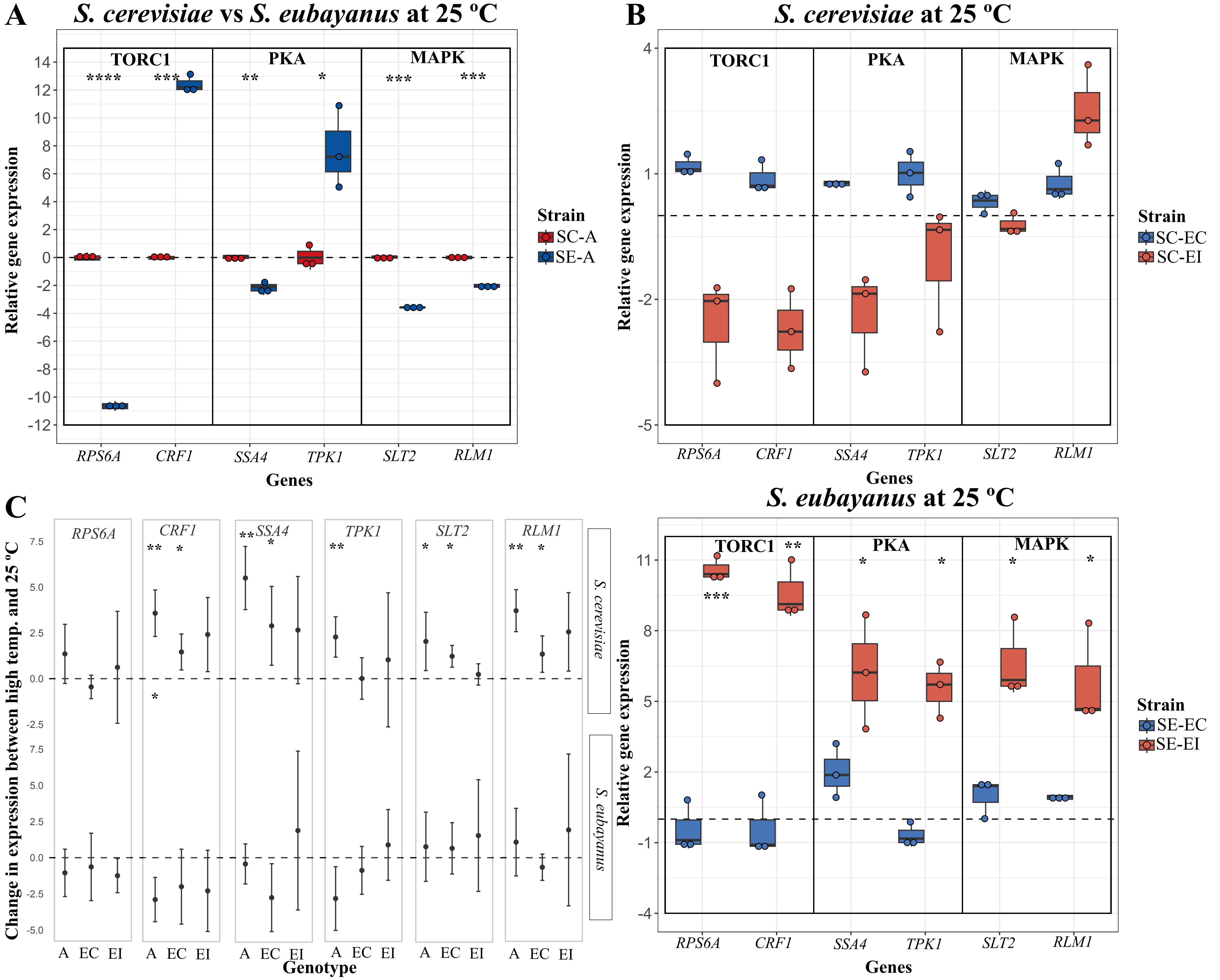


**Figure S2. Additional validation of TORC1/PKA/MAPK outputs across species and evolutionary conditions. (A)** Log2 normalized expression for ancestral *S. cerevisiae* (SC-A, red) and *S. eubayanus* (SE-A, blue) across the six genes at 25 ºC. Values are relative to SC-A. **(B)** Log2 normalized expression of evolved genotypes at 25 ºC relative to their species' ancestral strains at the same temperature (SC top; SE bottom). EC: genotype evolved at a constant 25 ºC; EI: genotype evolved under increasing temperatures. For (A) and (B), the dashed line marks no change. Plotted values correspond to the average of the biological replicates (n=3). Asterisks denote between-species differences in (A) and evolved-ancestral differences in (B) (Welch's t-test, * $p < 0.05$, ** $p < 0.01$, *** $p < 0.001$, **** $p < 0.0001$). Genes are grouped by pathway (TORC1: *RPS6A*, *CRF1*; PKA: *SSA4*, *TPK1*; MAPK: *SLT2*, *RLM1*). **(C)** Change in expression between each species' high temperature and 25 °C for A, EC, and EI (computed as high temperature minus 25 °C; high temperature is 40 °C for SC and 34 °C for SE). The horizontal dashed line marks no change. Asterisks denote differenceswithin genotype, per gene and species (Welch's t-test, * $p < 0.05$, ** $p < 0.01$, *** $p < 0.001$, high vs 25 °C ).
