## Supplementary material for "Convergent targeting of conserved regulatory networks during thermal evolution across *Saccharomyces*": Figure S3

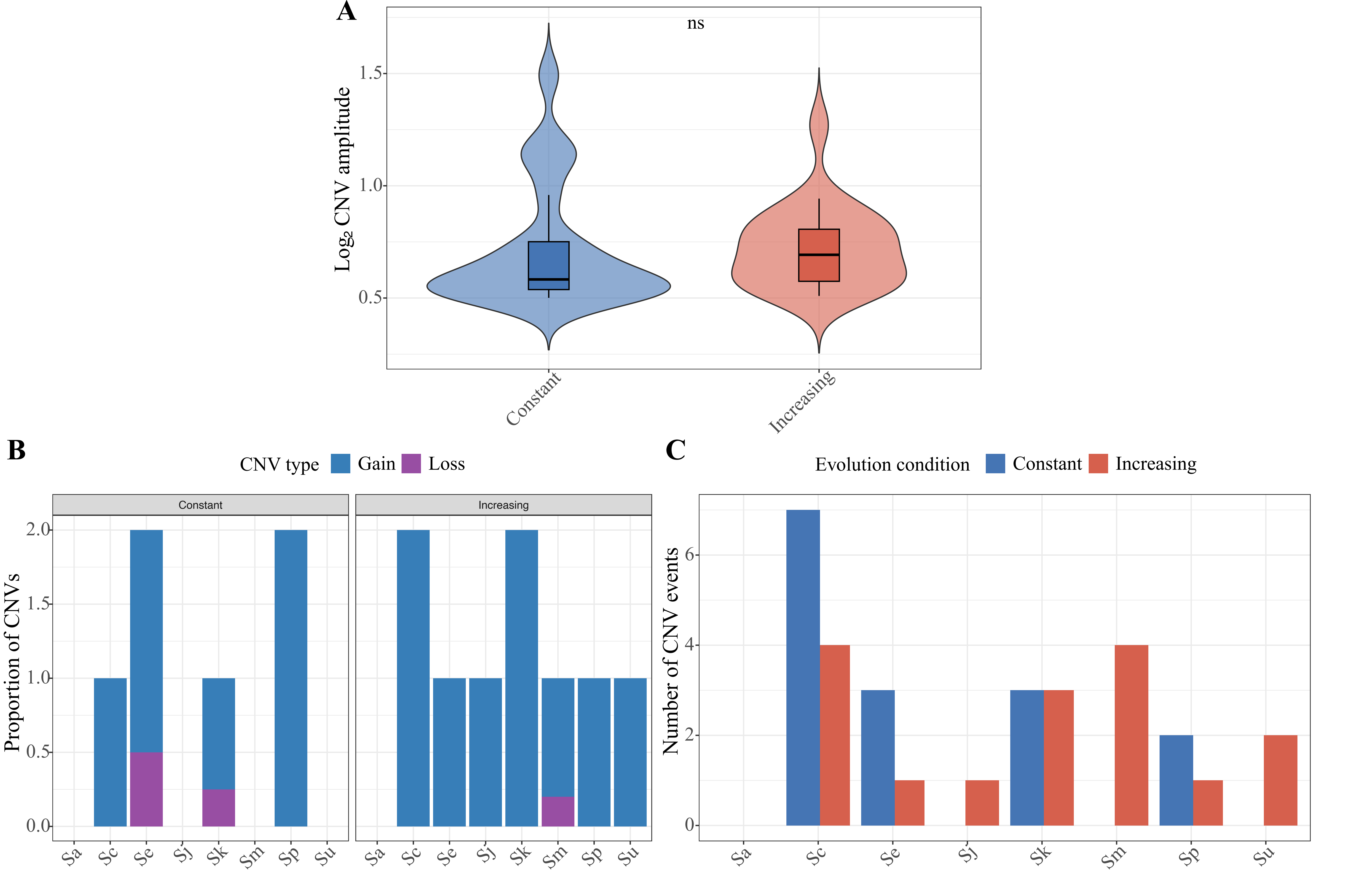


**Figure S3. Copy number variation associated with thermal evolution in *Saccharomyces*. (A)** Distribution of CNV amplitudes (absolute log2 copy-number change) for all significant CNV events detected across species under constant and increasing temperature evolution regimes. Violin plots show the density of CNV amplitudes, with embedded boxplots indicating median and interquartile range. CNV amplitudes did not differ significantly between evolutionary conditions (Wilcoxon rank-sum test, $p = 0.33$), indicating comparable magnitudes of copy-number changes across regimes. **(B)** Proportion of CNV gains and losses per species under constant and increasing temperature conditions. Stacked bars show the relative contribution of copy-number gains and losses within each species and condition, illustrating that gains were generally more frequent than losses across taxa. **(C)** Total number of CNV events per species under constant and increasing temperature conditions. Bars represent the number of condition-exclusive CNVs detected in each species, highlighting substantial species-specific variation but no consistent increase in CNV burden under increasing temperature evolution.
