## Supplementary material for "Convergent targeting of conserved regulatory networks during thermal evolution across *Saccharomyces*": Figure S4

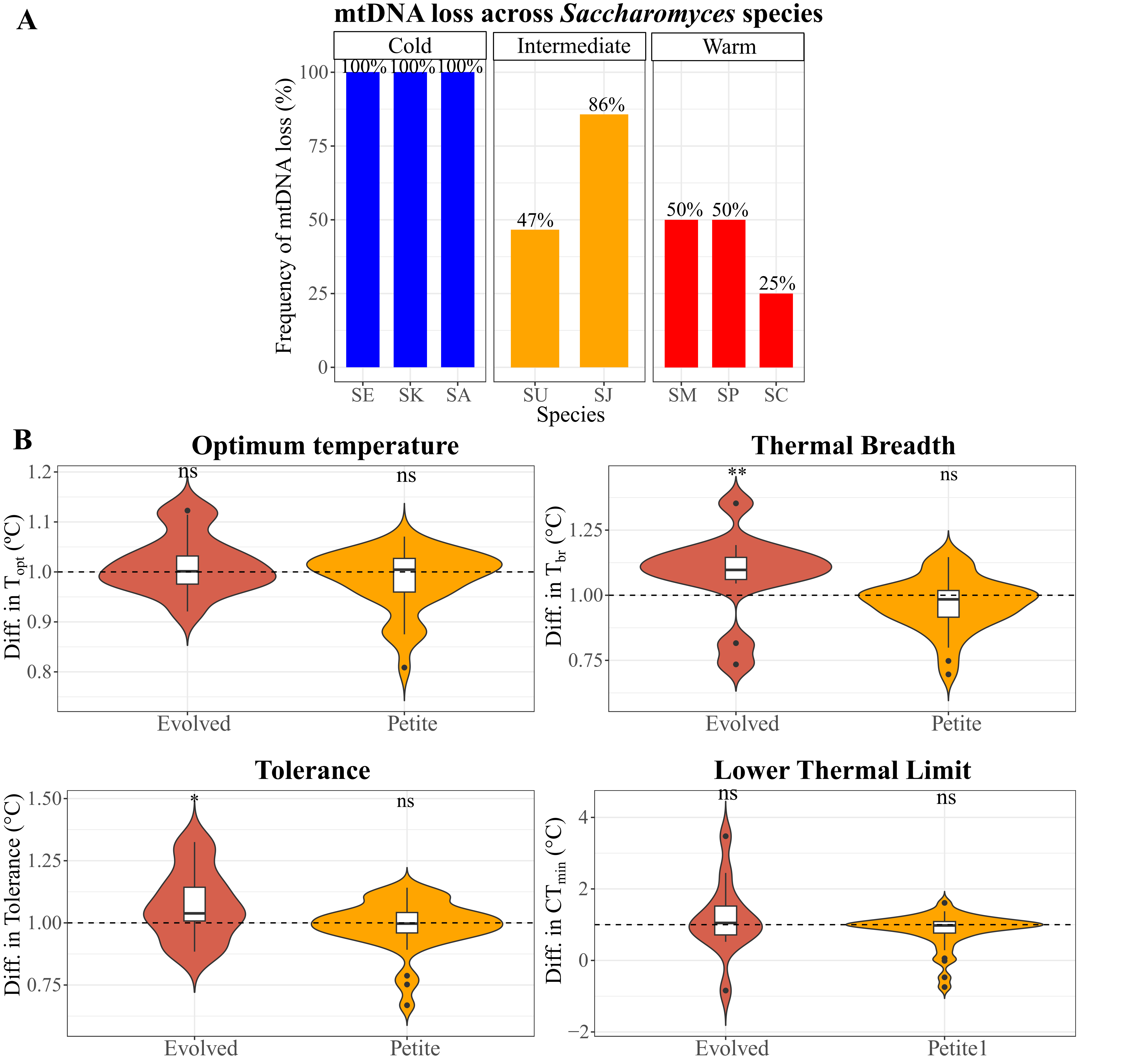


**Figure S4. Thermal parameters across ancestral, evolved, and petit strains.**
**(A)** Frequency of mitochondrial genome loss (petite formation) across *Saccharomyces* species. **(B)** Differences in thermal parameters between genotypes (Evolved vs Ancestral, Petite vs Ancestral). Violin plots show the distribution of differences in thermal parameters across strains, with internal boxplots indicating medians and interquartile ranges. The dashed line indicates the ancestral strains set at 1. Asterisks indicate significant differences between evolved and petit strains relative to their ancestors (Wilcoxon test; * $p < 0.05$, ** $p < 0.01$, ns not significant).
