## Supplementary material for "Convergent targeting of conserved regulatory networks during thermal evolution across *Saccharomyces*": Figure S5

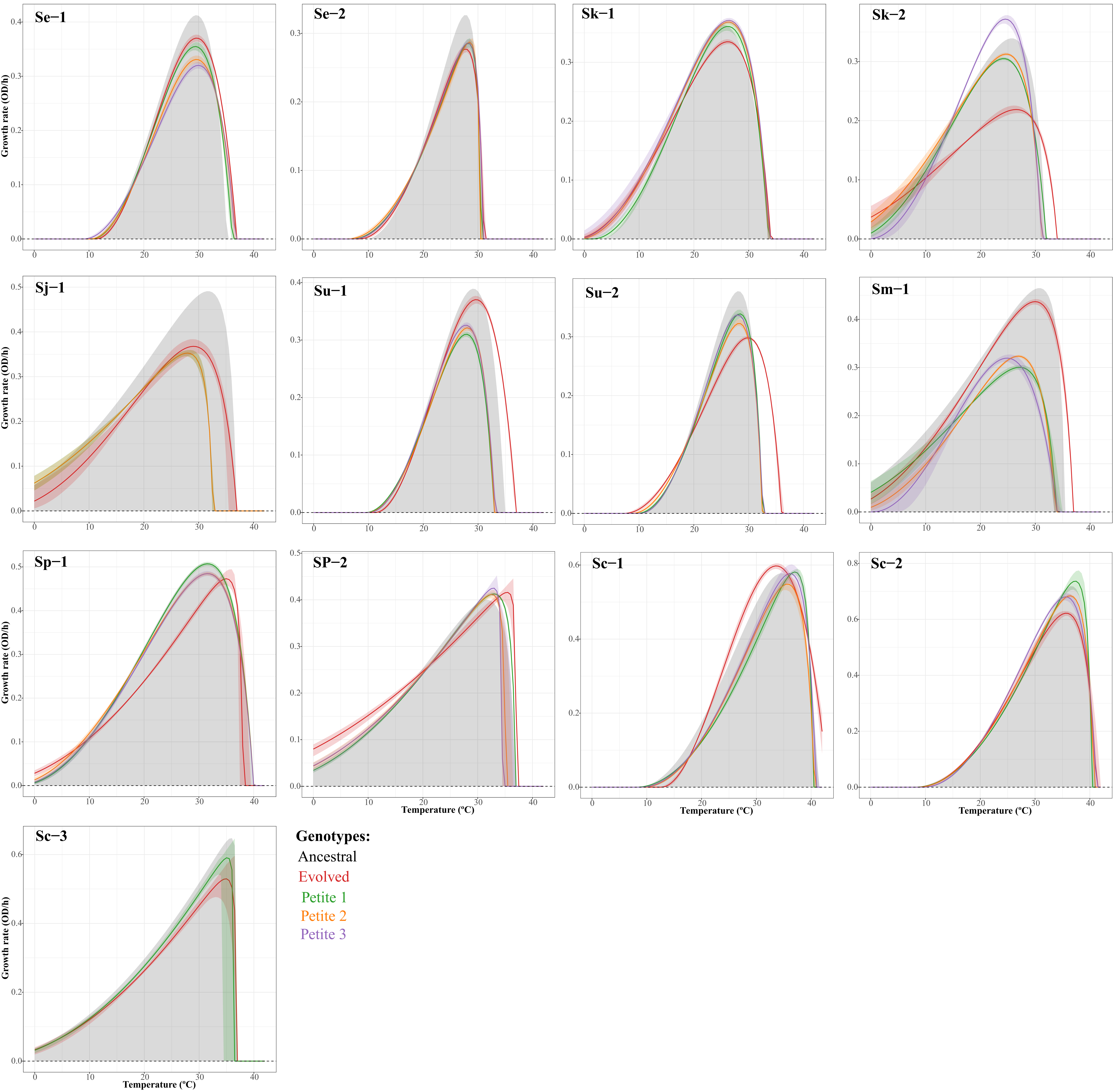


**Figure S5. TPCs of ancestral, evolved and petite genotypes per strain.**
Changes in maximum growth rates as a function of temperature for each strain, representing the TPCs. Ancestral TPCs are grey, evolved TPCs under increasing temperature are red, and petit TPCs are green, orange, and purple.
TPCs were constructed using the maximum growth rate of three technical replicates and were fitted using the CTMI.
